# Lysergic Acid Diethylamide extends lifespan in *Caenorhabditis elegans*

**DOI:** 10.1101/2025.06.16.659936

**Authors:** Beatriz de S. Carrilho, Aline Duarte, Isabelle Martins, Ariane Moura, Manuela Vilas, Matheus Antonio V. de C. Ventura, Francisco Moll, Hugo Aguilaniu, Ivan Domith, Stevens Rehen

**Author notes:** Corresponding authors at: D’Or Institute for Research and Education (IDOR), Rio de Janeiro 22281-100, Brazil. E-mail address (I. Domith); (S. Rehen).

## Abstract

Aging is modulated by nutrient-sensing pathways that integrate metabolic and hormonal cues to regulate growth, stress resilience, and lifespan. Caloric restriction (CR), a well-established intervention, extends longevity in diverse species through modulation of conserved nutrient-sensing pathways, including TOR signaling. Lysergic acid diethylamide (LSD), a classic serotonergic psychedelic with emerging therapeutic applications, remains largely unexplored in the context of aging. Here, we show that LSD treatment significantly extends lifespan in *Caenorhabditis elegans* and reduces age-associated lipofuscin accumulation, suggesting delayed age-associated decline. LSD induces several phenotypes that overlap with those observed in some caloric restriction paradigms, including reduced reproductive output, decreased body size, and the absence of an additive effect in lifespan assay performed in dietary restriction model, without apparent decreasing in food intake. In addition, LSD treatment modulates lipid stores and other cellular readouts associated with nutrient-sensing physiology. Together, these findings suggest that LSD engages evolutionarily conserved pathways linked to longevity and identify this compound as a tool to investigate serotonergic control of aging biology.

## Introduction

Aging is a complex biological process characterized by the progressive accumulation of sequential biological alterations over time, which increases susceptibility to disease and mortality (Harman, 1993). As global population ages, diseases such as cardiovascular conditions, diabetes, Alzheimer’s, and Parkinson’s disease have become increasingly prevalent, posing significant health and socioeconomic challenges (Fang et al., 2017). Addressing these challenges requires innovative strategies to extend longevity, delay the aging process, and mitigate the risk of age-associated diseases.

Over the past decades, research has focused on identifying and optimizing interventions to slow aging and promote healthier and longer lives (Hine et al., 2015; Jones et al., 2023; Kuhla et al., 2014). Among these, dietary modifications and regular physical activity have emerged as critical approaches. Specifically, calorie restriction (CR), defined as a reduction of 30–60% of caloric intake without malnutrition, stands out as one of the most effective strategies for extending lifespan and enhancing healthspan (Mercken et al., 2012). CR has been shown to significantly lower the incidence of aging-related diseases such as diabetes, cancer, heart diseases, and neurodegenerative disorders (Colman et al., 2009). Furthermore, fasting, which involves periodic abstinence from food, shares many benefits with CR, enhancing cellular resilience and reducing the risks associated with aging and disease (Longo & Mattson, 2014).

The mechanisms underlying the benefits of CR and fasting involve the activation of key molecular pathways, including the insulin/IGF-I-like (IIS) and the target of rapamycin (TOR) pathway. These signaling pathways play crucial roles in cellular protection against oxidative, metabolic, and proteotoxic stress, thereby enhancing longevity and delaying age-associated pathologies (Mattson, 2012). However, while the potential of CR to promote healthy aging is well-documented, sustaining a strict regimen in humans is challenging due to social, psychological, and practical constraints. This has led to growing interest in identifying alternative approaches, such as compounds that mimic the effects of CR without requiring drastic dietary restrictions.

Recent research has helped understand the interplay between dietary interventions, serotonin signaling, and aging mechanisms. Serotonin, traditionally recognized for its role in mood regulation, has emerged as a key regulator of aging and longevity (Fidalgo et al., 2013). Studies have shown that serotonin influences lifespan in several organisms, including *Caenorhabditis elegans* (Murakami & Murakami, 2007). In murine model, serotonin signaling is intricately linked to pathways such as mTORC1, which mediates the cognitive and metabolic benefits associated with dietary restriction (Teng et al., 2019). Targeting serotonin signaling may represent a novel strategy to promote longevity.

In mammals, serotonergic signaling can be modulated by psychedelics, which have gained significant attention for their potential to promote neuroplasticity, enhance mental health, and influence biological pathways linked to longevity (de Vos et al., 2021). Compounds such as LSD (Lysergic Acid Diethylamide), psilocybin, and N,N-dimethyltryptamine (N,N-DMT), are known to interact with serotonin receptors, impacting neuroplasticity and emotional regulation (Nichols, 2016).

Our recent findings highlight the use of *C. elegans* as a model organism for investigating the effects of psychedelics on serotonergic signaling and behavior. In this previous study, we demonstrated that acute LSD exposure reduces locomotion speed in *C. elegans* by 52%, mediated by serotonin receptors SER-1 and SER-4, which are homologous to human serotonin receptors 5HT2 and 5HT1, respectively (Ornelas et al., 2024). Additionally, metabolic profiling revealed that LSD is rapidly absorbed and metabolized into compounds such as Nor-LSD and O-H-LSD, analogous to metabolic pathways in humans (Silveira et al., 2023). While psychedelics have shown protective effects across various physiological systems, their role in modulating aging and promoting longevity remains an underexplored area of research (Inserra et al., 2023; Saeger & Olson, 2022). Given these findings, *C. elegans* emerges as an ideal model for further exploration of the link between serotonin signaling, psychedelics, and aging mechanisms. This study aims to investigate the effects of LSD on longevity. We hypothesize that LSD promotes longevity in *C. elegans* by modulating conserved nutrient-sensing pathways, potentially through mechanisms that overlap with aspects of caloric restriction physiology, thereby delaying physiological markers of aging.

## Methods

### Chemicals

Potassium phosphate monobasic (CAS 7778-77-0, Sigma Aldrich), Sodium phosphate dibasic (CAS 7558-79-4, Sigma Aldrich), Calcium chloride (CAS 10043-52-4, Merck), Magnesium sulfate (CAS 7487-88-9, Sigma Aldrich), Cholesterol (CAS 57-88-5, Sigma Aldrich), Streptomycin sulfate salt (CAS 3810-74-0, Sigma Aldrich), Agar bacteriological (Kasvi), Bacto peptone (Thermo Scientific), Potassium phosphate dibasic (CAS 7758-11-4, Thermo Scientific), LB Broth (Merck), Sodium chloride (CAS 7647-14-5, Merck), LSD (CAS 50-37-3, Lipomed), Oil Red O (CAS 1320-06-5, Sigma Aldrich), PBS (10X), pH 7.4 (Gibco), Triton^™^ X-100 (CAS 9036-19-5, Sigma Aldrich).

### Maintenance and cultivation of animals

The strains used in this study were obtained from the Caenorhabditis Genetic Center (CGC), University of Minnesota, USA. The strains N2 (Wild type), OH15876 [pha-4::3xGAS::GFP::TEV::LoxP::3xFLAG], AQ866 [(ser-4(ok512)], DA1814 [ser- 1(ok345)], DA1116 [eat-2(ad1116)], MAH240 [hlh-30p::hlh-30::GFP], and VC222 [raga-1(ok386)] were cultivated using standard techniques (Brenner, 1974). Briefly, the animals were maintained on solid medium in plates prepared with nematode growth medium (NGM) seeded with *Escherichia coli* OP50-1 as a food source and supplemented with 100 μg/L of streptomycin. The plates were kept at 20°C. To ensure adequate food supply, the animals were regularly transferred to fresh plates.

### Synchronization

The animals were synchronized using the bleaching method according to Porta-de-la Riva *et al*. (Porta-de-la-Riva et al., 2012). In summary, plates containing young adult hermaphrodites and eggs were collected and washed three times with M9 buffer to remove bacterial residues. Then, the bleaching solution (1.65% [v/v] NaCl, 0.5 M NaOH) was added, and the mixture was agitated to ensure complete lysis of the animals while preserving the eggs. M9 was added to stop the reaction, and the eggs were washed, transferred to bacteria-free plates, and stored at 20°C for 24 hours. The next day, L1 larvae were transferred to fresh plates containing bacteria to allow the animals to develop.

### Preparation for experiments

Except in the oviposition experiment, 5-fluorouracil was added to the medium at a concentration of 80 µM in long-term assays to prevent egg hatching. This compound interferes with DNA synthesis and blocks cell division. Additionally, to prevent the degradation of the compounds by bacteria, *E. coli* was previously inactivated by heat treatment at 65°C for 30 minutes.

### Longevity assay

Synchronized animals of the N2, raga-1(ok386), and eat-2(ad1116) strains at the young adult stage (day 0 of adulthood) were transferred to Petri plates containing either 250 nM LSD or vehicle. In the continuous exposure assay, animals remained on treatment plates throughout their lifespan, and survival was assessed every two days. Animals were considered dead if they showed no body or head movement upon mechanical stimulation. In parallel, a second longevity assay was performed to evaluate whether the longevity-promoting effects of LSD required continuous exposure. For this experiment, approximately 60 animals were allocated to each condition (Control or LSD 250 nM) and exposed to treatment for 24, 48, 72, 96, or 120 hours, after which both groups were transferred to treatment-free plates. Survival was then monitored every two days using the same scoring criteria. Survival analysis was performed using OASIS 2, and the results presented correspond to a late-survival time point, defined as the stage at which 25% of the animals remained alive.

### Fluorescence analysis and quantification

Fluorescence from the *pha-4::GFP* and *hlh-30::GFP* reporters were analyzed using the OH15876 and MAH240 strains, respectively, while lipofuscin autofluorescence was assessed in the N2 (wild-type) strain. Synchronized animals at the young adult stage (day 0 of adulthood) were placed on Petri plates containing either 250 nM LSD or not. Each plate corresponded to a specific time point: days 1, 5, 10 or 15 of adulthood. At the end of each treatment period, animals were imaged using a Leica M165 FC estereomicroscope at 120x magnification for *pha-4::GFP* and *hlh-30::GFP* fluorescence, and lipofuscin autofluorescence. Image analysis was performed using LAS X software. For *pha-4::GFP* and lipofuscin, total fluorescence was quantified in the head and in the whole body, respectively, and the fluorescence intensity values were normalized to the corresponding this region of interest (ROI) area. In MAH240 animals, HLH-30::GFP nuclear localization was assessed in intestinal cells based on the visual identification of GFP-enriched intestinal nuclei, and animals were classified as positive or negative for nuclear localization. Results were expressed as a percentage relative to the control group.

### Oviposition assessment

Synchronized adult animals were placed on Petri plates containing treatment (LSD 250 nM) or not. At the end of the L4 larval stage, animals were transferred to plates, with one animal per plate. Every 24 hours, the eggs were counted using a Leica Ivesta 3, and the adult animals were transferred to new plates containing the treatment or not, until day 10 of treatment.

### Animal body length

Synchronized animals at the young adult stage (day 0 of adulthood) were placed on plates containing either LSD (250 nM) or control conditions. Animals were assigned to plates corresponding to a single treatment time point (1 or 10 days). At the end of each treatment period, the animals were mounted on 2% agarose pads on microscope slides. Animal length was measured using LAS X software. Images were acquired using a Leica M165 FC stereomicroscope at 100× magnification.

### Pharyngeal pumping

Synchronized animals at the young adult stage (day 0 of adulthood) were placed on plates containing treatment (LSD 250 nM) or not. Approximately 12 animals were transferred to a plate containing 12 wells (one animal per well). Each well was recorded when the animal reached 1 and 5 days of treatment for 10 seconds using a Leica Investa 3 (Leica, Wetzlar, Germany) stereomicroscope with an attached camera and a 110x magnification. The analysis of each video was performed by playing them in slow motion (0.25×) and counting the pharyngeal pumping rate, with the counts independently assessed by two independent researchers to reduce individual observer bias.

### Lipid quantification

Synchronized animals at the young adult stage (day 0 of adulthood) were placed on Petri plates containing treatment (LSD 250 nM) or not. Approximately 100 animals were transferred to plates assigned to a single treatment time point (1, 5, or 10 days). Once the treatment ended, lipid quantification was performed according to Nicole Stuhr (Stuhr et al., 2022). In summary, the animals were washed, permeabilized with PBST, and fixed with 60% isopropanol. They were then incubated with Oil Red O 60% for staining. The animals were imaged using a Leica Ivesta 3 stereomicroscope 60x magnification (Leica, Wetzlar, Germany) with the same parameters for all experimental groups.

### Larval development

Synchronized animals at the L1 stage were placed on Petri plates containing treatment (LSD 250 nM) or not. Approximately 20 animals were transferred to each plate, and after 72 hours, the animals were scored for vulval development. The animals were imaged using a Leica SP8 confocal microscope.

### Statistical analysis

Lifespan assays were analyzed by comparing treated and untreated animals of the same strain using the log-rank (Mantel–Cox) test (See supplementary table 1). For all other experiments, statistical significance was assessed using the Student’s *t*-test or Two-way ANOVA, as indicated in the corresponding figure legends. In all experiments, each n was considered an independent experiment and were performed at least three times. All analyses were performed using GraphPad Prism version 8.0.

## Results

### LSD extends lifespan and reduces lipofuscin accumulation in aging *C. elegans*

Given that serotonergic signaling plays a crucial role in modulating longevity in *C. elegans* (Murakami & Murakami, 2007), we investigated whether chronic exposure to LSD increases nematode longevity. First, we performed a lifespan dose–response analysis to define the concentration used in subsequent experiments. Treatment with LSD at 100, 200, and 250 nM increased lifespan by 12.3%, 13.2%, and 17.2%, respectively, compared with controls (Supplementary Fig. 1). Since multiple doses were active in the nanomolar range, 250 nM was selected as the working concentration for subsequent experiments (Fig. 1a).

**Fig. 1.**
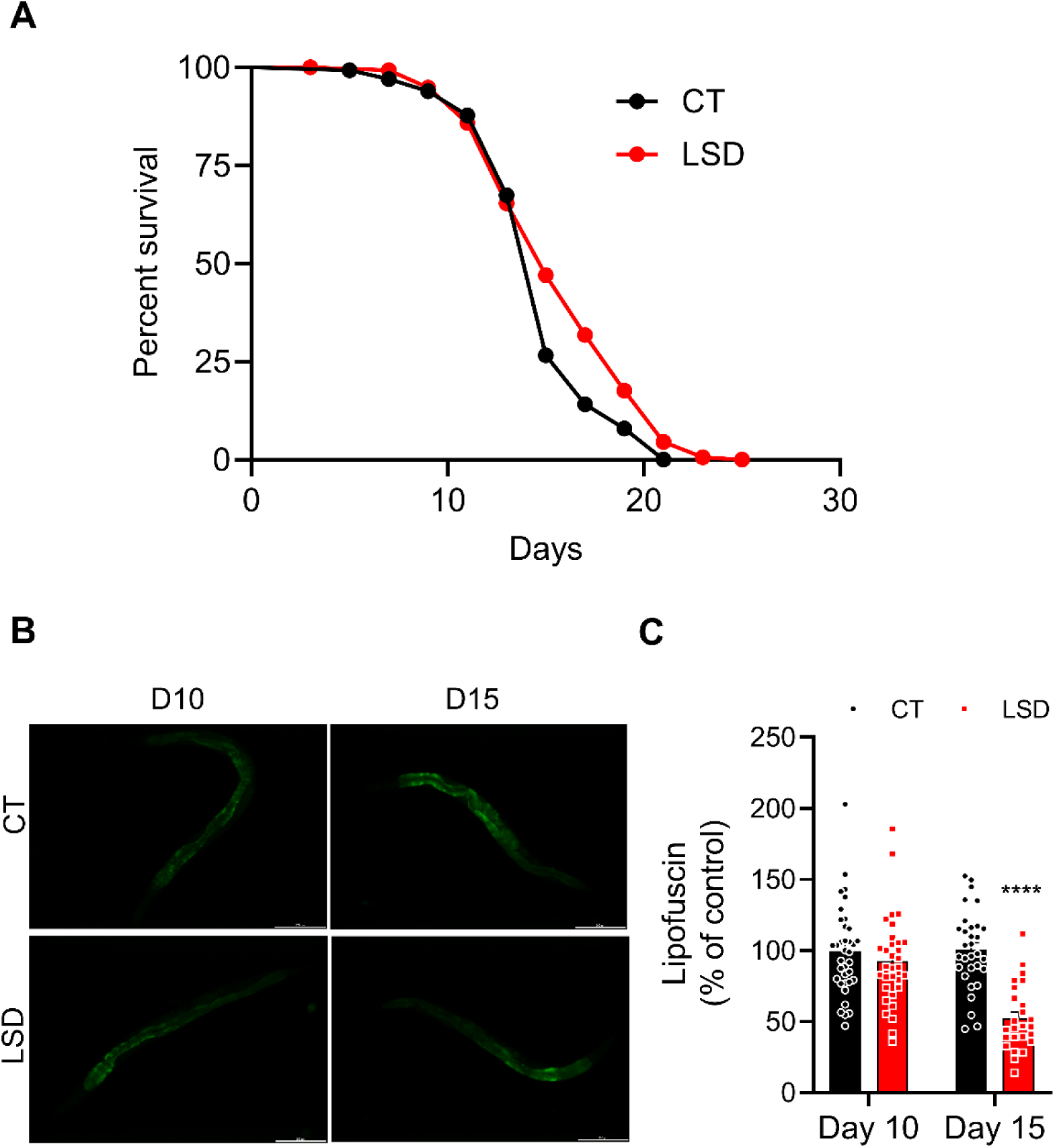
LSD extends the lifespan of *C. elegans* and delays aging. (**A)** Percentage survival of treated with 250 nM LSD (red) or untreated animals (black) over time. Data presented as Kaplan-Meier survival curves, and group differences analyzed using the Log-rank test. n=4, with at least 160 animals per condition. **(B)** Representative images of lipofuscin (green) animals treated (right) or untreated (left). Scale bar: 200 µm. **(C)** Quantification of lipofuscin accumulation in treated with 250 nM LSD (red) or untreated (black) animals on day 10 or 15. Fluorescence intensity expressed as percentage of the control and is presented as mean ± standard error of the mean (SEM). n=3, with at least 30 animals per condition. Statistical analysis performed using Student’s t-test. ****p<0.0001 compared to the control.

Although studying longevity provides insights into aging, increased lifespan does not necessarily reflect an extension of healthspan. To determine whether LSD also delays aging, we quantified lipofuscin, a fluorescent pigment that accumulates due to the incomplete degradation of cellular components. As its accumulation progressively increases with age, lipofuscin is considered a reliable marker of aging (Porta, 2002). We analyzed 10- and 15-day-old worms, as these time points allow sufficient lipofuscin accumulation to detect aging-related changes. LSD significantly reduced lipofuscin accumulation on day 15 compared with control, suggesting a delay in age-associated decline in addition to its lifespan-extending effect. (Day 10: CT 100% ± 10.5, LSD 100.7% ± 8.5; Day 15: CT 100% ± 6.4, LSD 49.8% ± 6.3, p < 0.001) (Fig. 1b and c).

### LSD-induced longevity depends on serotonergic signaling through SER-1 and SER-4

In our previous work, we identified SER-1 and SER-4 as key receptors involved in the behavioral changes triggered by LSD (Ornelas et al., 2024). Building on these findings, we investigated whether these receptors also contribute to the LSD-induced longevity. In *ser-4* mutants, LSD reduced lifespan, while in *ser-1* mutants LSD did not extend lifespan (Fig. 2a, 2b, and supplementary table 1). These results suggest that both receptors are necessary for LSD-mediated lifespan extension.

**Fig. 2.**
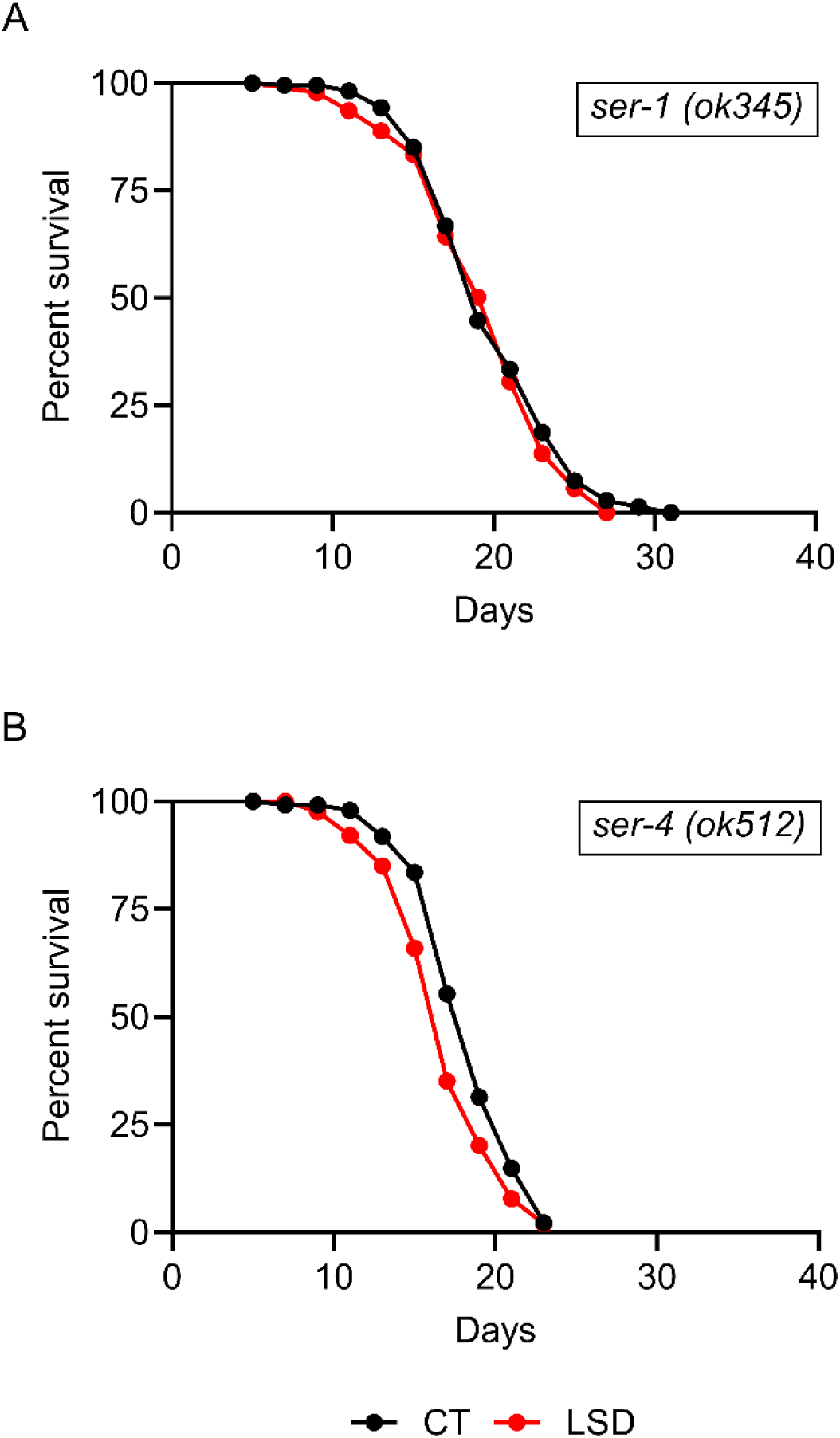
LSD promotes longevity through SER-1 and SER-4-dependent mechanisms. **(A)** Percentage of survival of mutant animals (*ser-1* KO) treated with 250 nM LSD (red) or untreated (black) over time. **(B)** Survival percentage of mutant animals (*ser-4* KO) treated with 250 nM LSD (red) or untreated (black) over time. All data presented as Kaplan-Meier survival curves, and group differences analyzed using the Log-rank test. n=4, with at least 160 animals per condition.

### LSD induces caloric restriction-like phenotype in *C. elegans*

In *C. elegans*, serotonergic signaling is not only involved in locomotor responses to food availability, such as the enhanced slowing response (Chase & Koelle, 2007), but also plays a broader role in nutrient-sensing and metabolic regulation. Since our previous work showed that LSD can elicit this response (Ornelas et al., 2024), we hypothesized that LSD-treated animals might display additional characteristics associated with CR.

A well-established consequence of CR in *C. elegans* is reduced oviposition (Hibshman et al., 2016). We quantified egg-laying in LSD-treated animals and observed that the effect of treatment varied over time (two-way ANOVA interaction, p<0.01; Fig. 3a), suggesting a CR-like modulation of reproductive output. CR is also known to reduce body size in *C. elegans* (Hibshman et al., 2016). We measured the length of treated and untreated animals on day 1 and 10 and identified that LSD treatment leads to a significant reduction in body length at all time points (Day 1: CT 849.5 µm ± 9.17, LSD 810.0 µm ± 10.27, p<0.01; Day 10: CT 1210.0 µm ± 15.86, LSD 1166.0 µm ± 12.37, p<0.05. Two-way ANOVA followed by Sidak’s multiple comparisons test) (Fig. 3c). A classical approach to induce dietary restriction in *C. elegans* is through genetic models with reduced pharyngeal pumping, such as the eat-2 (ad1116) strain. To assess the effect of LSD treatment in this context, we used this mutant in lifespan assays. No differences in survival were observed between control and treated animals (Fig. 3b). This result is consistent with a partial convergence between the effects of LSD and mechanisms associated with CR.

**Fig. 3.**
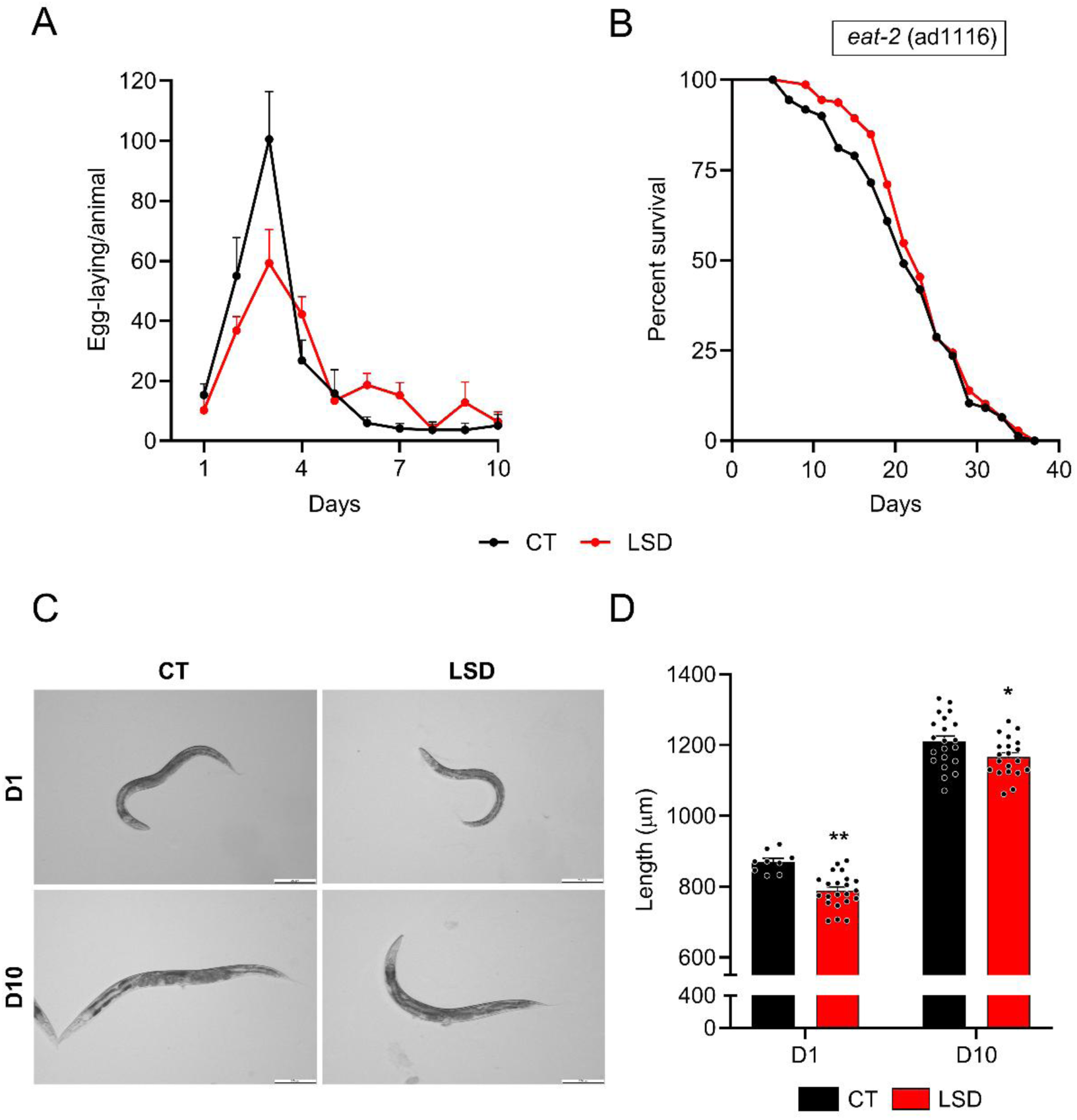
LSD induces caloric restriction-like phenotype. **(A)** Number of eggs laid during the reproductive period by treated animals with 250 nM LSD (red) or untreated animals (black) per animal/day. Data presented as mean ± standard error of the mean (SEM). n=3. Statistical analysis performed using Two-way ANOVA interaction, **p<0.01 compared to the control. **(B)** Percentage of survival of mutant animals (DA1116 *eat-2*) treated with LSD 250 nM (red) or untreated (black) over time. All data presented as Kaplan-Meier survival curves, and group differences analyzed using the Log-rank test. n=4, with at least 160 animals per condition. **(C)** Representative images of body length for treated (right) and untreated (left) animals over time. Scale bar: 200 µm. **(D)** Measurement of body length of treated animals with 250 nM LSD (red) or untreated animals (black) over time. Each point represents an individual animal. Statistical analysis was performed using two-way ANOVA followed by Sidak’s multiple comparisons test. *p<0.05 **p<0.01 compared to the control.

### LSD modulates caloric restriction-associated and metabolic outcomes over time without decreasing pharyngeal pumping

In wild-type animals, CR results from reduced food intake and is associated with adaptive changes that influence longevity. To assess whether LSD could indirectly trigger a CR-like response by reducing food intake, we measured the pharyngeal pumping rate of treated animals. Analysis on days 1 and 5 of adulthood revealed that LSD increased pharyngeal pumping frequency only on day 5 (Day 1: CT 229.0 ± 6.6, LSD 236.3 ± 7.6; Day 5: CT 185.0 ± 12.3, LSD 209.2 ± 12.2) (Fig. 4a). These findings argue against a simple reduction in food intake as the basis for the CR-like phenotypes observed previously and instead suggest that LSD engages aspects of nutrient-sensing physiology without suppressing feeding behavior under the conditions tested.

**Fig. 4.**
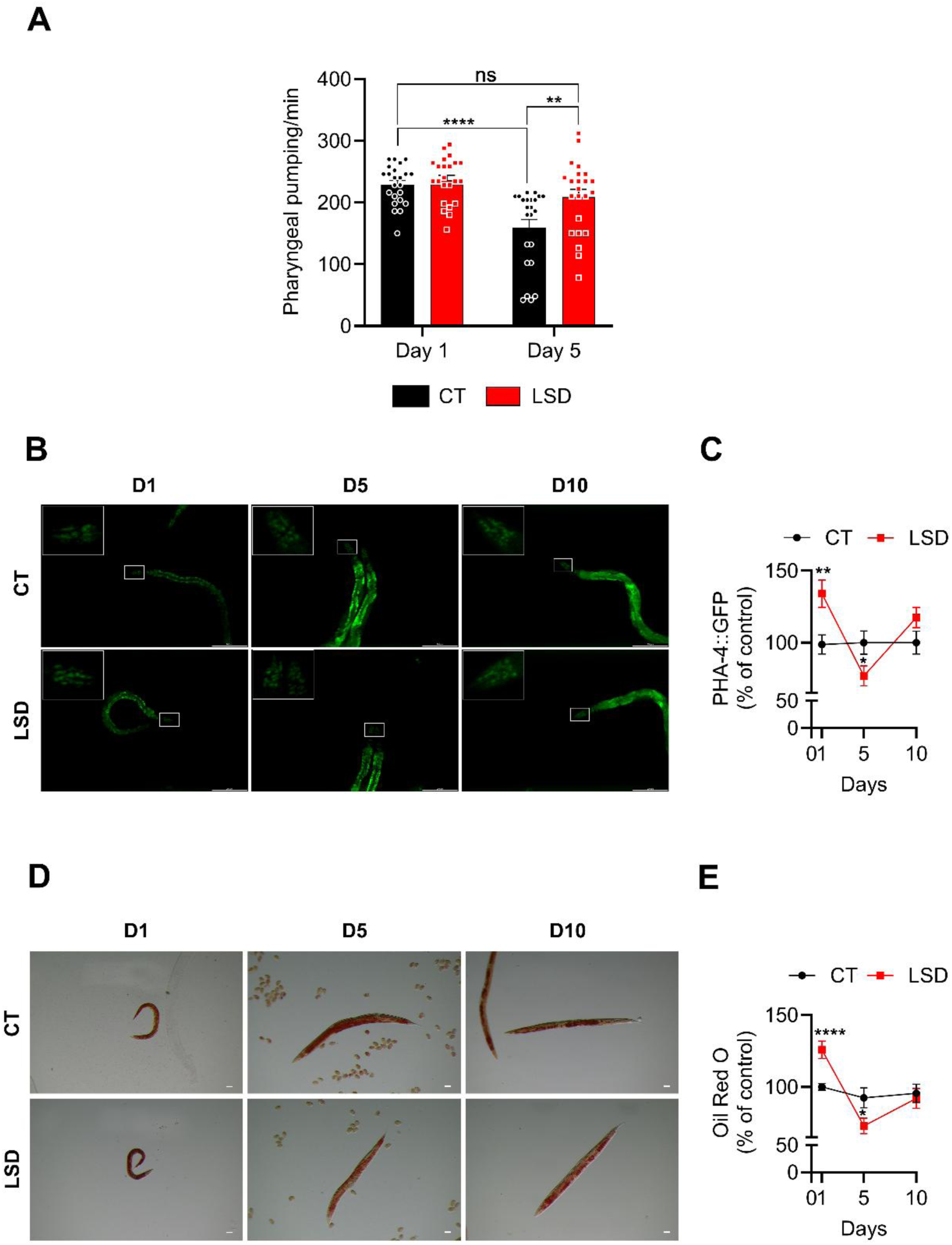
LSD modulates caloric restriction-associated and metabolic outcomes over time without decreasing pharyngeal pumping. **(A)** Quantification of pharyngeal pumping in treated with LSD 250 nM (red) or untreated animals (black) per minute. Each point represents an individual animal. Data presented as mean ± standard error of the mean (SEM). n=3. Statistical analysis was performed using two-way ANOVA followed by Sidak’s multiple comparisons test. **p<0.01; ****p<0.0001 compared to the control. **(B)** Representative images of untreated (left) or treated (right) PHA-4::GFP animals, with the inset in the upper left corner indicating the head region selected for fluorescence analysis. Scale bar: 200 µm **(C)** Quantification of PHA-4::GFP fluorescence intensity in the head region of treated (red) or untreated (black) animals. n=3 **(D)** Representative images of treated (right) and untreated (left) animals stained with Oil Red O (ORO) over time. Scale bar: 0.1 mm. **(E)** Quantification of total lipids in treated (red) or untreated animals (black) through ORO staining. n=3. **(B;D)** All data presented as mean ± standard error of the mean (SEM). Statistical analysis performed using Student’s t-test. *p<0.05; **p<0.01; ****p<0.0001 compared to the control.

Different dietary restriction paradigms can engage distinct cell signaling (Greer & Brunet, 2009). We therefore examined downstream readouts commonly associated with CR and nutrient-sensing physiology, including parameters linked to TOR signaling, and assessed whether LSD treatment modulates PHA-4 levels, a transcription factor acting downstream of TOR signaling, as well as lipid regulation. At day 1 of adulthood, treated animals displayed increased PHA-4 levels compared to control. This was followed by a reduction on day 5, while by day 10, levels were comparable to those of untreated animals (Day 1: CT 100.0 ± 6.5, LSD 134.2 ± 9.5, p<0.01; Day 5: CT 100.0 ± 8.11, LSD 76.9 ± 6.93 p<0.01; Day 10: CT 100.0 ± 7.96, LSD 117.4 ± 7.1) (Fig. 4b and 4c).

Similarly, lipid accumulation was dynamically regulated by LSD treatment. An increase in lipid content was observed on day 1, followed by a decrease on day 5, and a return to baseline levels by day 10 (Fig. 4d and 4e; Day 1: CT 100% ± 3.0, LSD 116.2% ± 6.6, p<0.01; Day 5: CT 90.3% ± 7.9, LSD 70.2% ± 5.8, p<0.01; Day 10: CT 93.1% ± 7.9, LSD 90.4% ± 10.1). Together, these findings reveal a transient and time-dependent metabolic remodeling in LSD-treated animals.

### LSD alters readouts commonly associated with nutrient-sensing pathways in *C. elegans*

To further explore cellular pathways that might contribute to the effects of LSD, we examined several developmental and transcriptional readouts previously linked to TOR pathway. As TOR signaling has been implicated in developmental timing, we assessed vulval formation in young adult nematodes to determine whether LSD induces a delay in larval growth (Jia et al., 2004). For this, we treated animals from the L1 stage and monitored continued until fully vulval maturation in young adults. Our results indicate that the number of animals with fully developed vulva was significantly lower in the LSD-treated group (CT 68.3% ± 5.3, LSD 31.7% ± 4.8, p<0.01) (Fig. 5a and 5b). This result is consistent with altered developmental timing and is compatible with modulation of TOR-related processes.

**Fig. 5.**
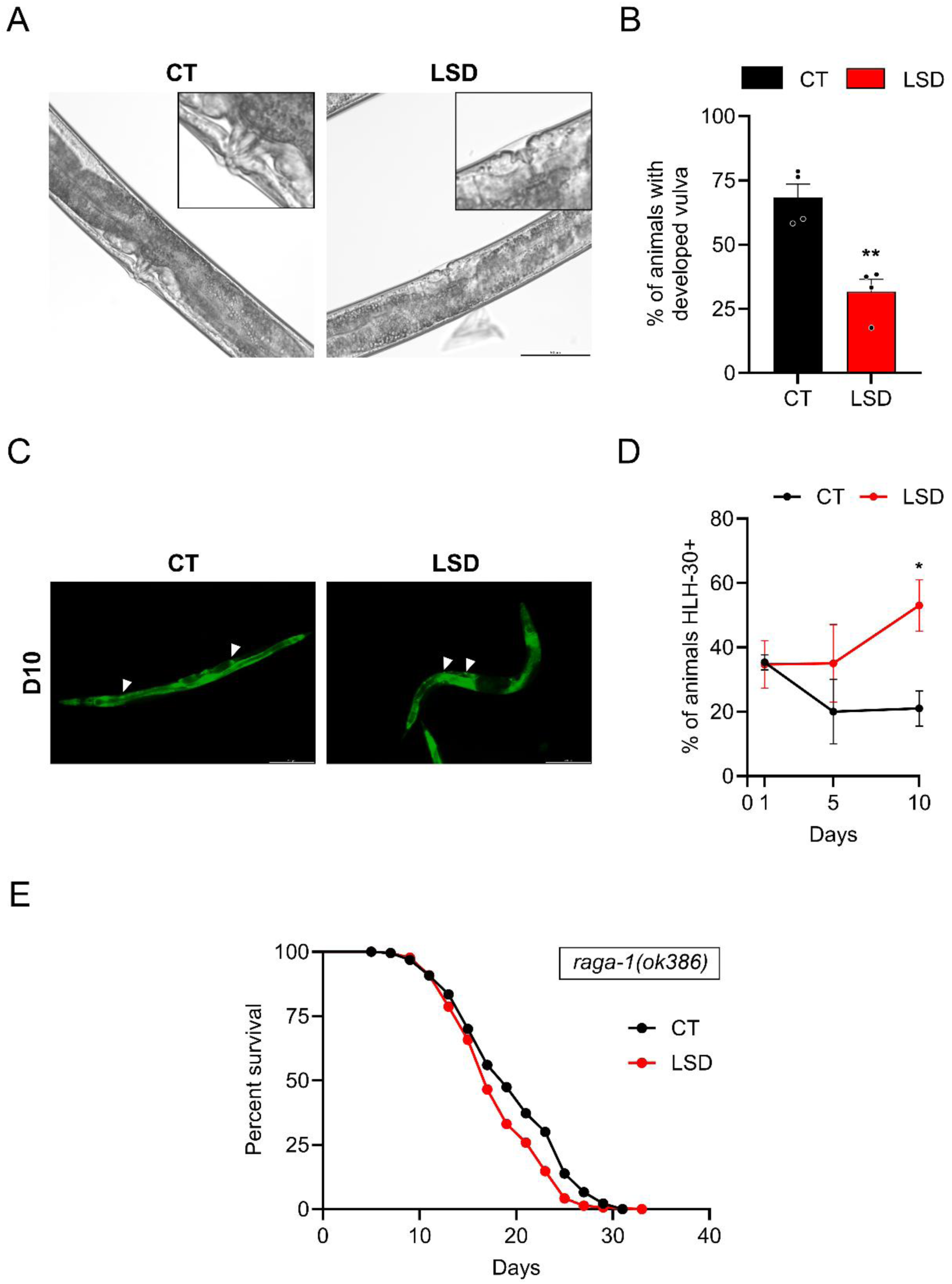
LSD affects developmental and transcriptional parameters associated with TOR signaling. **(A)** Representative image of the vulva (black arrow) in treated with LSD 250 nM (left) or untreated (right) animals. Scale bar: 50 µm. **(B)** Percentage of treated (red) or untreated (black) animals with developed vulvas at the end of the L4 larval stage. Each point represents an experimental n. n=4. **(C)** Representative images of HLH-30::GFP animals, treated (right) or untreated (left). Scale bar: 200 µm. **(D)** Quantification of animals displaying nuclear HLH-30::GFP localization in treated (red) or untreated animals (black). n=3. **(E)** Survival percentage of mutant animals (*raga-1* KO) treated with LSD 250 nM (red) or untreated (black) over time. All data presented as Kaplan-Meier survival curves, and group differences analyzed using the Log-rank test. n=4, with at least 160 animals per condition. **(B;D)** All data presented as mean ± standard error of the mean (SEM). Statistical analysis performed using Student’s t-test. *p<0.05; **p<0.01 compared to the control.

Next, we examined HLH-30/TFEB nuclear localization, as this transcription factor accumulates in the nucleus under conditions of reduced TOR activity (Silvestrini et al., 2018). LSD treatment increased the number of animals with nuclear translocation on day 10, but not on days 1 and 5 (Day 1: CT 35.3 ± 2.3, LSD 34.7 ± 7.4; Day 5: CT 20.0 ± 10.0, LSD 35.0 ± 12.1; Day 10: CT 21 ± 5.5, LSD 53.0 ± 8.0, p<0.01) (Fig. 5c and 5d). Finally, we performed a lifespan assay using the VC222 [*raga-1(ok386)*] strain, a deletion mutant affecting the Rag GTPase required for TORC1 activation. No differences in longevity were observed between control and LSD-treated animals (Fig. 5e). Together, these data are compatible with chronic LSD engaging TOR-associated processes, but do not by themselves demonstrate reduced TORC1 activity.

### LSD-induced longevity does not require continuous exposure

Because several LSD-induced physiological and metabolic changes were most evident during the first days of adulthood, we tested whether limiting exposure to this initial period would be sufficient to affect lifespan. Animals were therefore treated with 250 nM LSD for 24, 48, 72, 96, or 120 hours and then maintained in the absence of the compound for the remainder of life (Fig. 6). Under these conditions, all exposure time extended longevity (24 h: 5.6%; 48 h: 9.9%; 72 h: 11.0%; 96 h: 12.1%; 120 h: 11.0%). These results suggest that brief early-adulthood exposure can be sufficient to confer a longevity benefit; however, direct clearance controls are required to exclude residual exposure. Notably, the effect was detectable across multiple portions of the survival curve, suggesting that the benefit was not restricted to late survival alone.

**Fig. 6.**
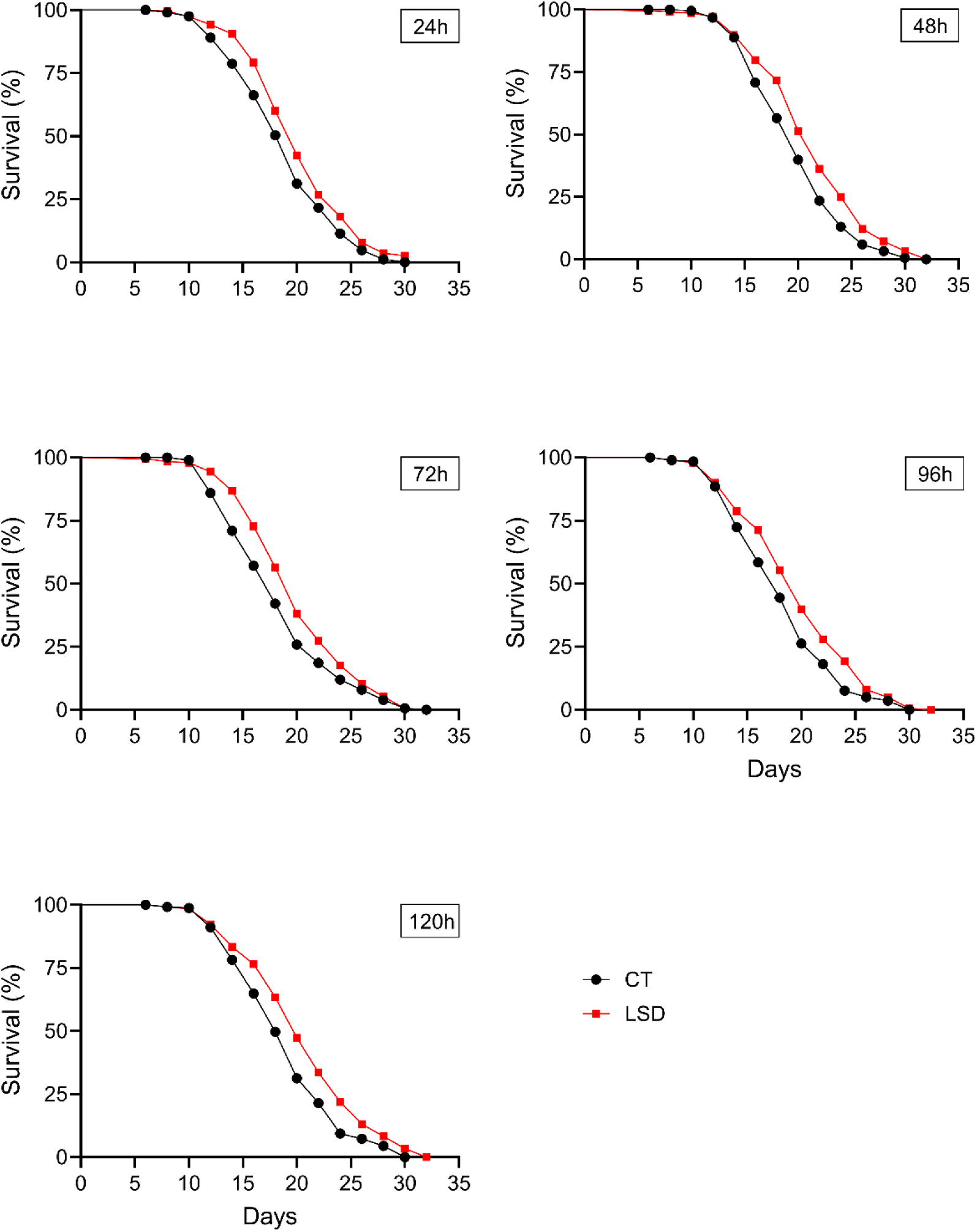
LSD treatment extends longevity independent of exposure duration. Percentage of survival of animals treated with LSD 250 nM for 24, 48, 72, 96 or 120 hours (red) or untreated (black) over time. All data presented as Kaplan-Meier survival curves, and group differences analyzed using the Log-rank test. n=3, with at least 180 animals per condition.

## Discussion

In this work, we demonstrate that LSD promotes lifespan extension while reducing lipofuscin accumulation, suggesting beneficial effects on both longevity and aging-related decline. These findings are consistent with an emerging view that psychedelics may influence core aging-related processes. Supporting this idea, a recent study reported that psilocybin/psilocin extended cellular lifespan in human fibroblasts and improved survival in aged mice, alongside reductions in senescence and oxidative stress-associated markers (Kato et al., 2025).

In our previous study, we showed that the behavioral slowing effect of LSD in *C. elegans* was absent in animals lacking serotonin receptors SER-1 and SER-4, indicating that both are involved in mediating its neuromodulatory actions (Ornelas et al., 2024). Building on those findings, we asked whether these receptors also play a role in the longevity effects induced by LSD. In *ser-1* mutants, LSD failed to extend lifespan, indicating that the pro-longevity effect of LSD is dependent on SER-1. In contrast, *ser-4* mutants showed a lifespan reduction upon LSD treatment. This suggests that, in the absence of SER-4, LSD may either exert a deleterious effect or counteract the lifespan increase typically observed in *ser-4* mutants. Altogether, these results indicate that LSD-induced longevity is receptor-specific and depends on the organization of serotonergic signaling rather than on a generic serotonergic effect.

The longevity effects of LSD exhibit parallels with those of serotonergic transporter inhibitors such as fluoxetine, paroxetine, and sertraline, which extend lifespan in *C. elegans* by modulating serotonin receptor pathways (Liang et al., 2024; C. Zhou et al., 2024; Y. Zhou et al., 2024). Notably, the serotonin antagonist mianserin has been proposed to mimic CR, a phenomenon wherein lifespan extension is achieved without reducing nutrient intake, by engaging serotonin receptor-mediated signaling (Petrascheck et al., 2007). Similarly, LSD may share some of these effects, as our results suggest the modulation of features compatible with CR, including reduced egg laying (Hughes et al., 2007) and body size (Hibshman et al., 2016), as well as the absence of an additive effect in lifespan assays using the *eat-2* strain. Taken together, these findings are consistent with partial overlap between the effects of LSD and physiological programs associated with CR. Importantly, this pattern does not appear to result from reduced food intake. On the contrary, we observed that, by day 5 of adulthood, treated animals exhibited increased pharyngeal pumping activity. As this activity naturally declines with aging (Cho et al., 2025), maintenance of higher pumping rates at day 5 is also compatible with delayed physiological decline, in agreement with the reduction in lipofuscin accumulation.

In many dietary restriction paradigms in *C. elegans*, increased longevity is accompanied by lipid metabolic remodeling and, frequently, by a reduction in fat content during adulthood. However, this response is not uniform and may vary depending on the protocol, the developmental stage analyzed, and the severity of the restriction (Palgunow et al., 2012). Indeed, different CR interventions engage distinct metabolic pathways, and in some contexts, dietary restriction can even lead to a transient increase in triglyceride levels and expansion of lipid droplets (Greer & Brunet, 2009). In this context, the increase in lipid content observed after 1 day of LSD treatment may reflect an early phase of metabolic reorganization. The reversal of this trend by day 5, when lipid levels decrease, suggests that the treatment induces a dynamic and time-dependent response consistent with a progressive adaptation of energy metabolism.

In parallel, PHA-4 expression appeared to follow a similar pattern, with an increase on the first day of treatment followed by a reduction on the fifth day. Previous studies have demonstrated the essential role of this transcription factor in lifespan extension associated with dietary restriction, involving both its nuclear activity and increased expression (Panowski et al., 2007; Sheaffer et al., 2008). In addition, PHA-4 has been shown to promote lipid accumulation under conditions of nucleolar stress through the transcriptional activation of lipogenic genes (Wu et al., 2018). In this context, the increase in PHA-4 observed after 1 day of LSD treatment is compatible with an early response that overlaps with nutrient-sensing (Panowski et al., 2007) or anabolic-state remodeling, although these data do not distinguish between these possibilities (Sheaffer et al., 2008), even in the absence of a detectable reduction in food intake. At the same time, this early increase in PHA-4 may also help explain the transient rise in lipid content observed at day 1, consistent with a phase of metabolic reorganization (Wu et al., 2018). In turn, the decrease observed on day 5 may indicate that this initial response is not sustained at the same level over time, which is consistent with a transient physiological adaptation (Cornwell et al., 2024), followed by the establishment of a new state of equilibrium.

Although LSD exhibits features consistent with a CR-like response, its effects may not fully overlap with those of classical CR mimetic agents. For example, rapamycin, which also modulates nutrient-sensing pathways via mTOR inhibition, is associated with adverse effects such as glucose intolerance and increased diabetes risk (Unnikrishnan et al., 2020). Psychedelics, in contrast, have been associated with reduced metabolic risks, such as protective effects on pancreatic β-cells, lower odds of diabetes and heart disease s (Gojani et al., 2024; Simonsson et al., 2021). However, these observations should not be taken as direct evidence that LSD reproduces the full physiological profile of CR mimetics. In the context of the present study, the main conclusion is more limited: LSD induces a physiological program in *C. elegans* that shares several features with CR-associated responses.

Furthermore, TOR pathway plays a pivotal role in growth, metabolism, and aging (Keith Blackwell et al., 2019). In our study, LSD delayed larval development, reduced body size and increase of HLH-30 nuclear translocation at later time point. Together, these observations are compatible with modulation of TOR-associated processes, as described in dietary restriction and rapamycin paradigms (Bar et al., 2016; Duong et al., 2020; Keith Blackwell et al., 2019). Psychedelics like LSD and DMT activate 5-HT2A receptors, which have been shown to mediate mTOR activation during short-term treatments, such as those enhancing neuroplasticity in cortical neurons (Ly et al., 2018; N. Costa et al., 2024). Proteomic analyses of overnight LSD-treated human cerebral organoids reveal upregulation of proteins involved in mTOR signaling and synaptic plasticity, further supporting this mechanism during acute exposure (N. Costa et al., 2024). Whether prolonged LSD exposure produces signaling adaptations distinct from acute exposure remains to be determined. Studies on 5-HT6 receptor signaling demonstrate that receptor desensitization and decreased membrane expression attenuate mTORC1 activity over time, a phenomenon that may also apply to 5-HT2A receptors with prolonged LSD exposure. Our data are consistent with a temporal shift in TOR-related readouts, but they do not establish direct TOR modulation as the sole mechanism underlying the longevity phenotype.

When we investigated whether the longevity-promoting effects of LSD required continuous exposure, we found that shorter treatment regimens (72, 96, and 120 hours) each produced a similar beneficial effect. Together, these results indicate that LSD can induce long-lasting physiological changes sufficient to promote lifespan extension even after limited exposure, a feature that may be especially promising in a therapeutic context.

## Conclusion

In conclusion, our findings demonstrate that LSD extends lifespan and attenuates age-associated physiological decline in *C. elegans*. This effect depends, at least in part, on serotonergic signaling and is accompanied by physiological and metabolic changes consistent with modulation of conserved nutrient-sensing pathways. Rather than acting through reduced food intake, LSD appears to induce a time-dependent adaptive response associated with longevity-related biology. Together, these results position LSD as a useful tool to investigate how serotonergic signaling interfaces with aging-related pathways in *C. elegans*.

## Acknowledgements

We would like to thank Dr. Marcelo Alves da Silva Mori for reviewing the manuscript and for helping with the acquisition of mutant animals. The graphical abstract was created with BioRender.

## Funding sources

This work was partially supported by the intramural grant from D’Or Institute for Research and Education, and IDOR/ Pioneer Science Initiative (www.pioneerscience.org). Students’ scholarships were provided by CAPES and IDOR.

## Declaration of competing interest

The authors declare that they have no known competing financial interests or personal relationships that could have appeared to influence the work reported in this paper.

## CRediT authorship contribution statement

**Beatriz de S. Carrilho:** Writing – original draft, Methodology, Investigation, Formal analysis, Conceptualization. **Aline Duarte:** Writing– review & editing, Methodology, Investigation. **Isabelle Martins:** Methodology, Investigation. **Ariane Moura:** Methodology, Investigation. **Manuela Vilas:** Methodology, Investigation. **Matheus Antonio V. de C. Ventura:** Methodology, Investigation. **Francisco Moll:** Investigation. **Hugo Aguilaniu:** Writing – review & editing, Supervision. **Ivan Domith:** Writing – original draft, Validation, Investigation, Formal analysis, Supervision. **Stevens Rehen:** Writing – review & editing, Supervision, Resources, Conceptualization, Funding acquisition. Overall, artificial intelligence was used to support language revision and improve textual clarity.

## Notes

### Competing Interest Statement

The authors have declared no competing interest.

### Summary of Updates

In this revised version, we repeated the experiments using a different concentration and updated the analyses based on these new results. We also added new experiments to further investigate and clarify the mechanism responsible for the observed effect. These additions improve the robustness of the findings and refine the interpretation of the study. The manuscript has been updated throughout to reflect these changes in the Results, Methods, Figures, and Discussion.

