## Supplemental material for "Lysergic Acid Diethylamide extends lifespan in *Caenorhabditis elegans*"

**Table 1 – Effects of LSD on lifespan**

| Strain | Treatment time | LSD concentration (nM) | Number of animal (LSD-/LSD+) | Number of experiments | Percentage change | P value |
|---|---|---|---|---|---|---|
| **N2 (Wild Type)** | Whole life | 100 | 137/174 | 4 | 12.4 | 0.0321 |
| **N2 (Wild Type)** | Whole life | 200 | 137/174 | 4 | 13.2 | 0.013 |
| **N2 (Wild Type)** | Whole life | 250 | 137/271 | 4 | 17.6 | 0.0026 |
| **N2 (Wild Type)** | 24h | 250 | 215/240 | 4 | 5.6 | 0.0027 |
| **N2 (Wild Type)** | 48h | 250 | 231/219 | 4 | 9.9 | 0.0007 |
| **N2 (Wild Type)** | 72h | 250 | 202/199 | 4 | 11 | 0.0063 |
| **N2 (Wild Type)** | 96h | 250 | 206/209 | 4 | 12.1 | 0.0035 |
| **N2 (Wild Type)** | 120h | 250 | 210/238 | 4 | 11.1 | 0.0001 |
| **ser-4 (ok512)** | Whole life | 250 | 267/212 | 4 | 7.7* | 0.0001 |
| **ser-1 (ok345)** | Whole life | 250 | 240/230 | 4 | 2.2* | 0.358 |
| **raga-1 (ok386)** | Whole life | 250 | 224/230 | 4 | 11.6* | 0.023 |
| **eat-2 (ad1116)** | Whole life | 250 | 131/140 | 4 | 3.5 | 0.398 |

**Supplemental table 1. Summary of lifespan assays in *C. elegans*.** Lifespan assays were performed in wild-type (N2), ser-1(ok345), ser-4(ok512), raga-1 (ok386) and eat-2 (ad1116). Worms were exposed to 250 nM LSD either chronically (whole-life) or for defined durations (24–120 h). The number of animals analyzed per group (control or LSD-treated), number of biological replicates, and percentage change in 75% lifespan relative to controls are shown. Statistical significance was determined by comparing survival curves of treated and untreated animals of the same strain using the log-rank (Mantel–Cox) test. * For these strains, the percentage difference indicates that lifespan was higher in control animals than in LSD-treated animals

**Supplemental figure 1.**

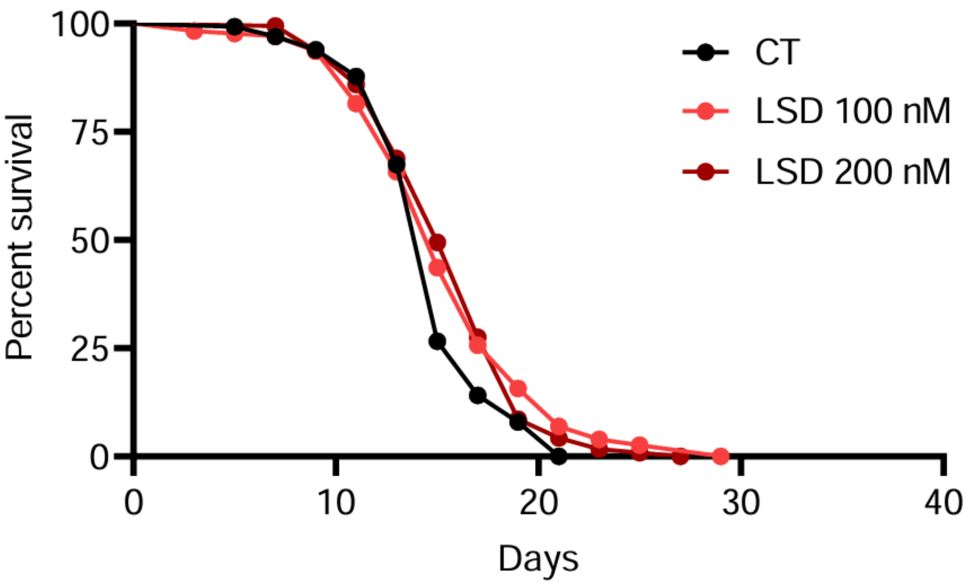


**Supplemental figure 1**. **Lifespan response to different LSD concentrations**. Percentage survival of treated with 100 or 200 nM LSD (orange and red, respectively) or untreated animals (black) over time. Data presented as Kaplan-Meier survival curves, and group differences analyzed using the Log-rank test. n=4, with at least 160 animals per condition.
